# Subcellular Region Morphology Reflects Cellular Identity

**DOI:** 10.1101/2024.08.14.607889

**Authors:** Ángel-Carlos Román, Alba Diaz-Pizarro, Nuria Del Valle-Del Pino, Marcos Olivera-Gómez, Guadalupe Cumplido-Laso, Dixan Agustín Benítez, Jose María Carvajal-González, Sonia Mulero-Navarro

## Abstract

In multicellular organisms, various cells perform distinct physiological and structural roles. Traditionally, cell identity has been defined through morphological features and molecular markers, but these methods have limitations. Our study explores the potential of subcellular morphology to define cellular identity and predict molecular differences. We developed workflows to identify subcellular regions in different cell lines, using convolutional neural networks (CNNs) to classify these regions and finally quantify morphological distances between cell types. First, we demonstrated that subcellular regions could accurately distinguish between isolated cell lines and predict cell types in mixed cultures. We extended this approach to predict molecular differences by training networks to identify human dermal fibroblast subtypes and correlating morphological features with gene expression profiles. Further, we tested pharmacological treatments to induce controlled morphological changes, validating our approach in order to detect these changes. Our results showed that subcellular morphology could be a robust indicator of cellular identity and molecular characteristics. We observed that features learned by networks to distinguish specific cell types could be generalized to quantify distances between other cell types. Networks focusing on different subcellular regions (nucleus, cytosol, membrane) revealed distinct morphological features correlating with specific molecular changes. This study underscores the potential of combining imaging and AI-based methodologies to enhance cell classification without relying on markers or destructive sampling. By quantifying morphological distances, we provide a quantitative characterization of cell subtypes and states, offering valuable insights for regenerative medicine and other biomedical fields.

## Introduction

In multicellular organisms, various cells perform distinct physiological and structural roles. Depending on the species, organ, or tissue, multiple cell types and subtypes have been described from different points of view (1, 2). Since the 19th century, we have understood that the cell serves as the fundamental unit of life, with each cell originating from another (3). Despite the possibility of identical genomic content among different cells within an organism, several molecular mechanisms drive variations in their morphology and functionality (4–6). Additionally, the same cell type can exist in different states based on environmental stressors (7, 8). Regenerative medicine aims not only to characterize cells but also to convert one cell type into another, with cell reprogramming and differentiation as central concepts in the field (9–11).

But how do we define the identity of a cell? (12) Ideally, we would classify each cell based on its function (13). As a direct proxy for cell function, during most of the previous century, microscopists were able to classify distinct cells by their subcellular morphology and their ability to be stained by chemical molecules (14), with the main limitation being their visual acuity. With the advancement of molecular biology, several proteins have been defined as cell type markers for use in immunofluorescence (15), flow cytometry (16), or *in situ* RNA hybridization experiments (17). Implementing this strategy across all cells is challenging due to the need to know the cell markers in advance and the existence of cell subtypes with minor differences between them (18). Beyond cellular function, obtaining transcriptomic profiles of single cells (scRNA-seq) has overcome the limitations of protein markers (8, 13, 19, 20). Different algorithms can classify known cells and even identify new cell types (21). Additionally, scRNA-seq data theoretically allows us to extract molecular distances between cells to define subtypes, and recent advances are bringing us closer to obtaining spatial information in scRNA-seq (22). What are the main limitations of transcriptomic profiling of single cells? Apart from the elevated cost, the major issue with scRNA-seq is the required destruction of the cellular sample during RNA extraction, which hinders subsequent analyses or therapies involving selected cell types or subtypes (23).

In summary, there are several alternative approaches to characterizing cellular identity from distinct morphological or molecular perspectives. From a functional standpoint, morphological features of the cell are closer to its identity than molecular profiles. One reason for this is the increased variability of transcriptomes, which can lead to errors in identifying cellular types or states. Our hypothesis is that subcellular regions contain sufficient information to distinguish between different cellular identities. For instance, micro-shapes of the cellular or nuclear membrane, as well as the number and size of organelles, can serve as relevant features for classifying different cell types (12, 13). By modifying classic microscopy-based strategies using imaging and AI-based methodologies, we can enhance their effectiveness and classify cell types without relying on markers or causing sample destruction. Additionally, we can estimate morphological distances between cells, quantitatively characterizing various cell subtypes or states. Finally, these morphological distances can be evaluated for their similarity to molecular differences among cell types. In our study, we demonstrate that identifying and comparing subcellular regions across different cell types allows us to determine their cellular identity. Furthermore, these subcellular morphological distances can predict their molecular counterparts. Remarkably, this adaptable model can be applied to multiple theoretical or practical cellular contexts.

## Results

To test our hypothesis that subcellular information can predict cell identity, we first designed a general workflow for identifying subcellular regions from different cellular samples. In its simplest form (Figure 1A), it classifies a subcellular region as one of two distinct cell types, such as the human breast cancer cell lines MCF7 or MDA-MB-231. The process involves the following steps: First, we obtained a library of images from white light microscopy of both cell lines. We used a 10x objective to retrieve between 20 and 40 images per cell line. Second, a simple transformation to black and white from grayscale followed by a contrast thresholding algorithm and a windowing function generated smaller Regions of Interest (ROIs) containing subcellular regions from the initial images. For all the experiments shown here, we obtained between 40,000 and 100,000 subcellular ROIs. Third, the two subcellular ROI libraries were used as input for a standard CNN that learned to classify a subcellular region between both cell lines. After training, we were able to predict whether an unknown subcellular region came from MCF7 or MDA-MB-231 (Figure 1A). Using a non-previously trained dataset of cellular images, we quantified the accuracy of cell type classification. As a proof of principle, we assessed the accuracy of one-by-one classifications of twelve different cell lines (Figure 1B). The accuracy of the prediction was high in all cases, with the lowest being 856‰ for Phoenix vs. NTERA cells. The best identification accuracy was for HL60, which produced an accuracy over 985‰ in all cases, supporting our hypothesis since HL60 is a semi-adherent cell line with morphological properties very different from the other cell lines (Figure 1B). We also tested the system’s capability by generating CNNs that learned to distinguish between four or eight different cell lines (Figure 1C). The accuracy of prediction dropped to around 700‰ with eight cell lines, while it remained around 900‰ with four cell lines, suggesting that the current workflow can handle four cell lines simultaneously. Given that cellular confluence is a relevant parameter in cell cultures, we tested its effect on cell line identification. We typically used highly confluent (70%-90%) cell images to minimize the number of initial pictures needed. Then, using a CNN trained to identify HEK293 or MDCK under these conditions, we predicted its accuracy with lower confluent cellular images (Figure 1D). We found that the prediction accuracy was not affected by reduced cell confluency (over 990‰). A potential drawback of this workflow is the need for isolated images of the different cell types to be recognized. We addressed this by demonstrating its application in a mixed cell culture (Figure 1E). Specifically, we combined unmarked HEK293 with marked (live cell red marker) MDCK cells and obtained images in both white light and the red channel. We modified the workflow from Figure 1A to obtain subcellular ROIs only from regions positive in the red channel. Using these labeled ROIs, we achieved accuracies similar to those obtained with the standard method (isolated, unmarked cells) (Figure 1E).

**Figure 1.**
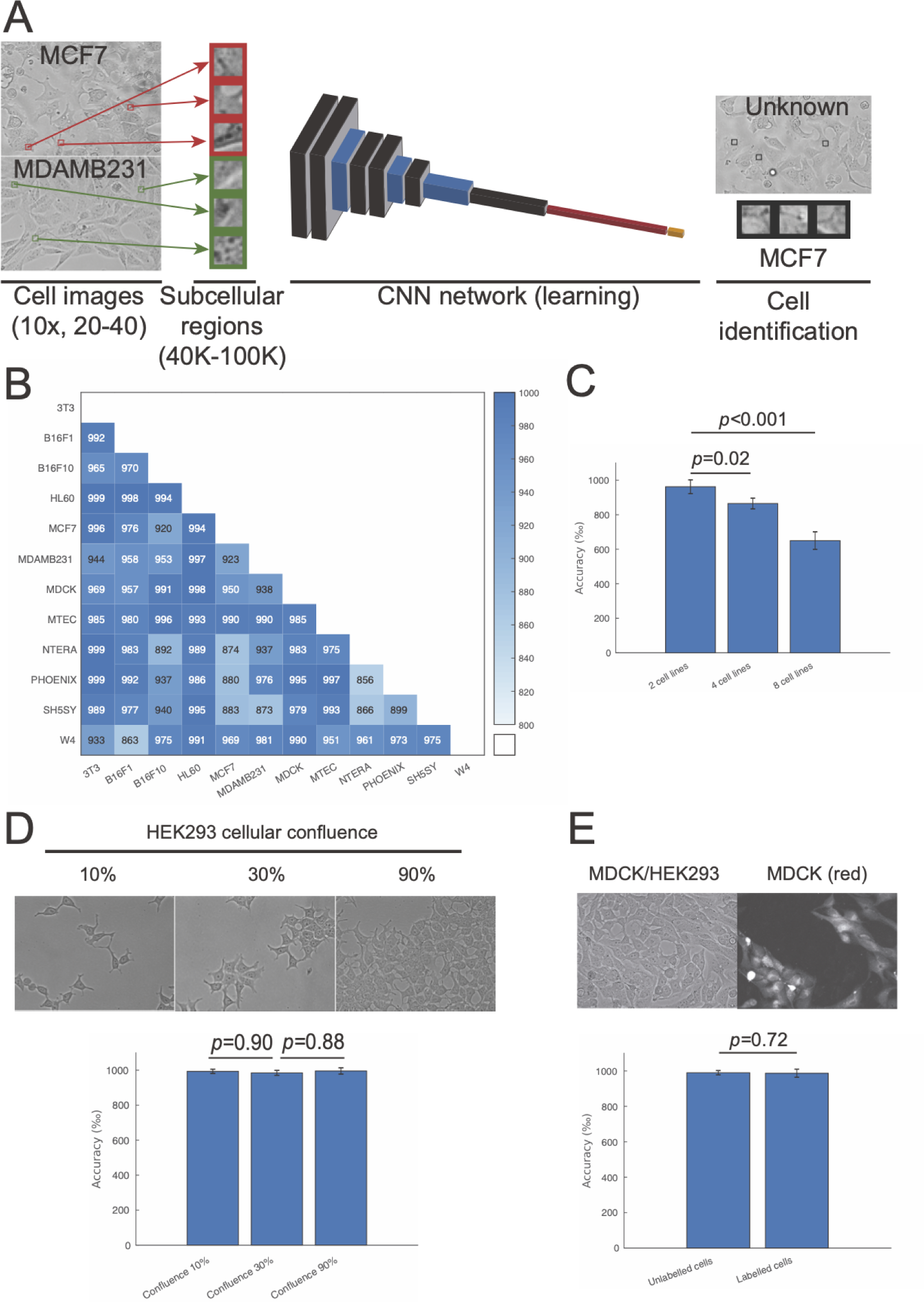
Subcellular regions can classify different cell lines. (A) Scheme of the CNNs that are used along the manuscript. (B) Triangular matrix indicating pairwise accuracy in the detection of cell lines. (C) Average accuracy of CNNs classifying two, four or eight cell lines at the same time. (D) Accuracies in the detection of cell lines at different levels of confluence. (E) Comparison of accuracy with single cell culture or the use of labelled cells in co-culture as input for CNN.

So far, we have observed that subcellular region morphologies contain enough information to discriminate between different isolated cell lines. However, in many situations, several cell types are mixed in the same culture, as shown in Figure 1E. We wanted to evaluate the accuracy of our method in the case of a co-culture of two different cell lines (HEK293 and MDCK, Figure 2). In addition to white light images (Figure 2A and A’), HEK293 cells were live-cell marked (Figure 2B and B’) to identify them and apply an automatic grayscale threshold to calculate the positive pixels (Figure 2C and C’). A modified version of our method scans the image for ROIs identified as either HEK293 or MDCK and then generates a prediction in the image (Figure 2D and D’). Finally, the accuracy of the direct threshold and the prediction was estimated by comparison to a gold standard (human-based calculation using the white light and live-cell marker images, Figure 2E and E’). We used the real images (Figure 2A-E) as well as a magnification (Figure 2A’-E’), obtaining good predictive results in both cases, with real-size images generating better results than the magnifications (Figure 2E and E’). We tested whether these predictions could be used to estimate the amount of a specific cell type present in a co-culture. We prepared different co-cultures composed of controlled percentages of four cell lines (MCF7, MDA-MB-231, MDCK, and B16F1) and used a CNN trained to distinguish between these cells to predict the percentage of each cell line in each co-culture (Figure 2F). To optimize the system, we divided the number of pixels of each cell line by an estimation of the pixel size of that cell line (see Materials and Methods for more details). We observed a strong correlation between predicted and real percentages of cell types (Figure 2F). As a final assessment of the efficacy of subcellular region morphologies for determining cell type within co-cultures, we tested whether dynamic changes in cell morphology could be detected using our method. For this, we obtained images during a trypsin-based MDCK cell detachment process. We used the initial MDCK images (before trypsin) and the final images of the process (when MDCK cells are rounded) to create a CNN that can distinguish between both states. Then, we predicted the percentage of MDCK in each state observed during the detachment (Figure 2G), successfully reconstructing the detachment dynamics of MDCK from subcellular region morphologies.

**Figure 2.**
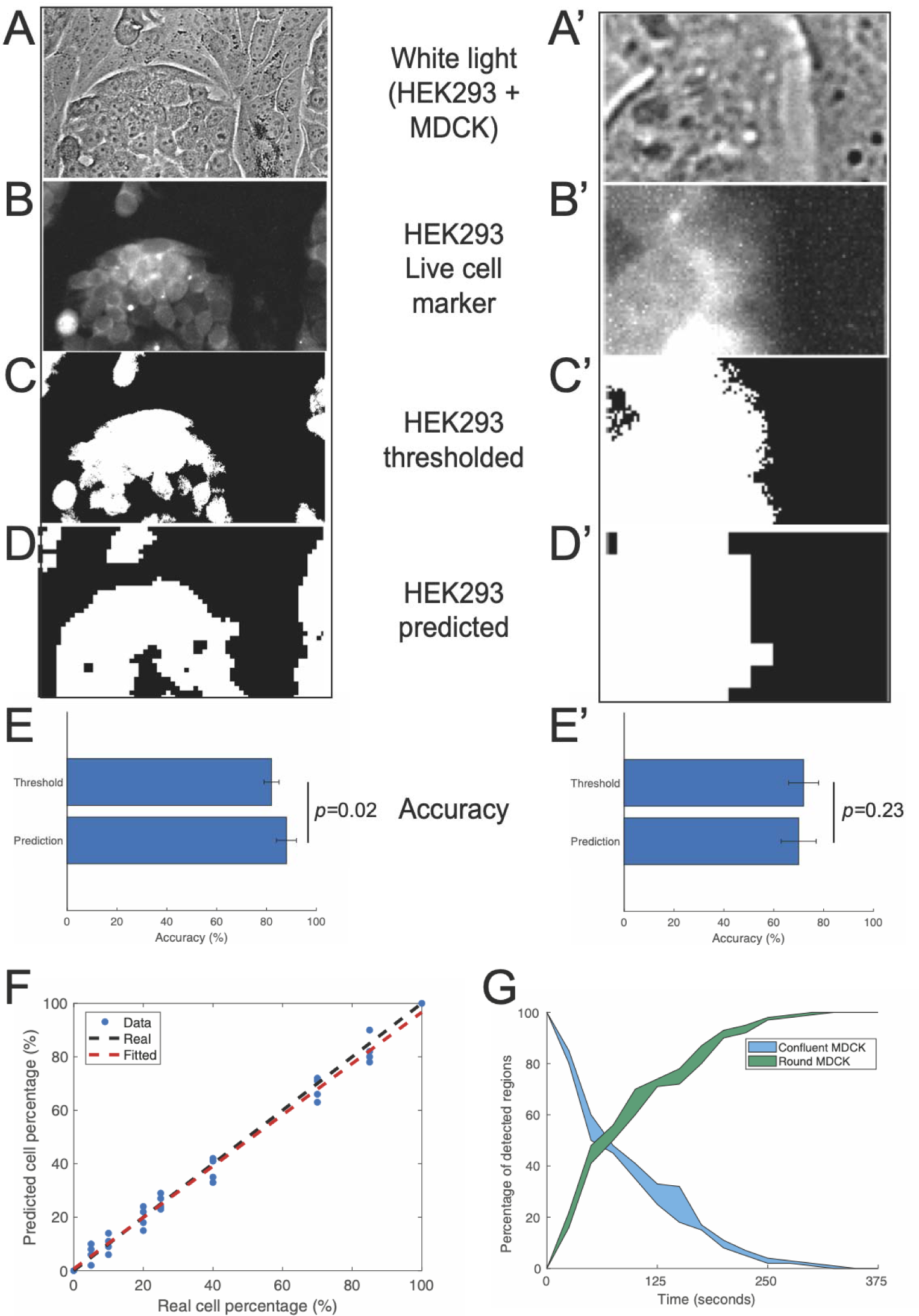
Applications of cell classification using CNNs. (A-D, A’-D’) Two examples of *in situ* detection of cell lines in a co-culture (left, original image; right, a magnification of another region), from (A-A’) white light images, the (B-B’) labelled cells under the red channel, the application of a (C-C’) contrast threshold in this channel and the (D-D’) prediction of the CNN. (E-E’) Quantification of accuracy in the detection using the prediction and the contrast threshold of the marker, compared to the human definition of the cell contours. (F) Predicted vs. Expected percentage of different cell lines within a co-culture, with a fitted line from the results. (G) Dynamic quantification of MDCK subtypes (confluent vs. round) along the detachment process.

Next, we assessed whether the information contained in cellular morphologies could be used to predict molecular differences between cellular subtypes. For example, human papillary and reticular dermal fibroblasts are characterized by differential expression of gene markers such as UCP2 (in papillary dermal fibroblasts) and COL11A1 (in reticular dermal fibroblasts) (24). These dermal subtypes have different properties regarding the cell microenvironment, among other factors (25). This is relevant due to the extensive use of dermal fibroblasts from patient skin biopsies for human Induced Pluripotent Stem cell (hIPS) cell reprogramming. We built a network that learned to distinguish between subcellular regions from human papillary and reticular dermal fibroblasts. Then, we determined the subtype (papillary/reticular) of several clones from commercial human dermal fibroblasts as well as from a donor skin biopsy (Figure 3A and B). Additionally, we quantified the gene expression of papillary (UCP2, Figure 3A) and reticular (COL11A1, Figure 3B) markers in these clones. We observed a strong correlation between morphology and molecular predictions in the case of human dermal cell subtypes, as papillary or reticular marker expression is consistent with a papillary or reticular morphology, respectively. This led us to further investigate the ability of subcellular region morphology to quantify subtle changes between distinct cells. To do this, we established a culture of proximal airway basal stem cells from adult mice tracheas (MTBSCs) (26). We developed a network that could identify MTBSCs or HEK293 regions, and then used this network to quantify its accuracy using not only the original MTBSCs cell sample but also other MTBSCs obtained from other mice (MTBSC2 and MTBSC3, Figure 3C). Additionally, we cultured all the MTBSCs for four additional passages and repeated the accuracy prediction using the original network (Figure 3C). We observed that MTBSCs subcellular regions seem to be more similar between different cell culture batches than between different culture passages (Figure 3C). However, it is difficult to assign small changes in accuracy to differences in cell morphology, so we developed a simple method to generate cellular distances from the network (see Materials and Methods). For each region analyzed, we extracted a feature vector from the last pooling layer; then, we combined the set of regions from the same sample using dimensionality reduction followed by covariance analysis to obtain a representation of each sample in the Euclidean space (Figure 3D). This confirmed the previous result (Figure 3C) regarding the similarity between different batches of MTBSCs and the differences observed after several cell passages.

**Figure 3.**
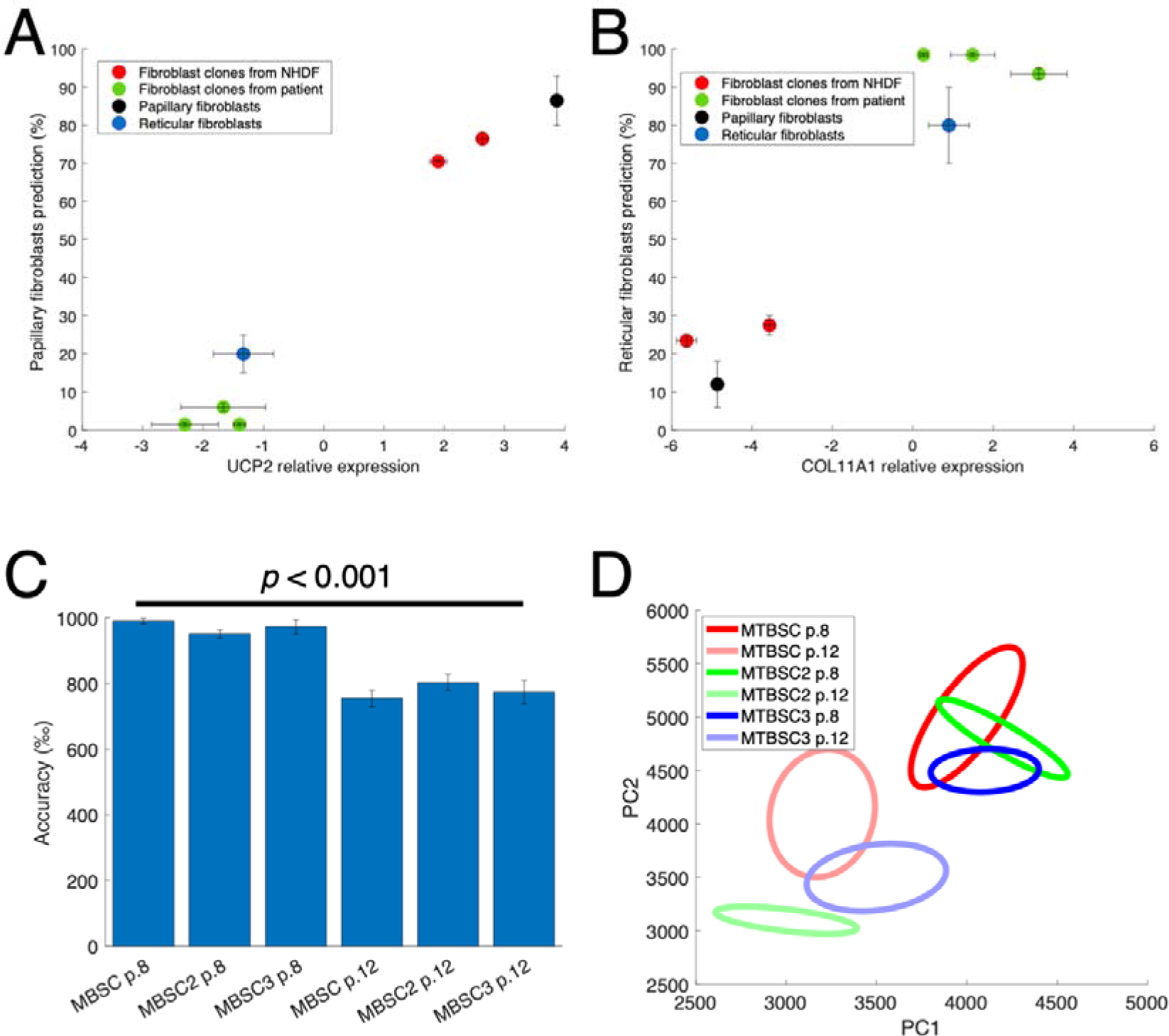
Morphological distances between cell types and comparison to molecular markers. (A) Quantification of relative expression of UCP2 gene (papillary marker) compared to estimation of cell type using a CNN. (B) Same as in A, but using the relative expression of COL11A1 (reticular marker). (C) Quantification of accuracy in the detection of different batches or passages of MTBSCs. (D) Visualization of morphological distances between batches or passages of MTBSCs using principal component analysis (PCA) followed by estimation of ellipse rotation and size using the eigenvectors.

Now we have shown that we can quantify morphological distances between cell types using subcellular features learned from the original samples. Nevertheless, using cellular features obtained from other classification networks that do not include the original cell types is theoretically feasible. To validate this idea, we compared the distances between MTBSCs batches obtained using the initial network (MTBSC vs. HEK293) with three other networks (MTBSC vs. MCF7, MCF7 vs. HEK293, and MDCK vs. HEK293), two of which do not include MTBSCs (Figure 4A, see Materials and Methods for the explanation of distance quantification). We observed a strong positive correlation between the original distance and those obtained using other networks (R>0.9, p<0.001). This result indicates that the morphological features learned to distinguish specific cell types can be generally used for distance quantification between other distinct cell types. This led us to further explore the possibility of generating alternative networks for distance quantification. We used a similar approach as in Figure 1E (live nuclei markers) to generate four different networks that included different subcellular regions of HEK293 and MDCK cell lines. Regions located less than 15 pixels from the nuclear signal (15px network), between 15 and 30 pixels (30px network), between 30 and 45 pixels (45px network), and between 45 and 60 pixels (60px network). First, we quantified the accuracy of these networks to detect potentially more informative subcellular regions within the cell (Figure 4B). Nuclear and membrane (15px and 60px) regions seemed to be slightly more informative than cytosolic regions (30px and 45px), but there were no significant differences among them. Then, in parallel to Figure 4A, we quantified the distance between a set of six cell lines (HEK293, MCF7, MDA-MB-231, HEPG2, HL-60, and SH5-SY5Y) using the 15px, 30px, 45px, and 60px networks as well as another network that did not include subcellular fragmentation (Figure 4C). We found that the distances obtained varied depending on the type of network used. There was a strong correlation between 15px and 30px distances (R=0.89, p<0.001) and between 30px and 45px distances (R=0.64, p=0.01), but in the rest of the pairwise comparisons, the correlation was non-significant (R<0.47, p>0.05). This result suggests that these networks are learning different features for cellular classification, related to the different positioning of the regions used, from nuclei to membrane. We then wondered if these distinct features could be related to specific molecular changes between these cell lines, as molecular profiles are also considered a strong indicator of cell identity (13). To address this hypothesis, we analyzed the public transcriptomic data of these cell lines in the Cancer Cell Line Encyclopedia (CCLE) (27), see Materials and Methods for a detailed explanation of the process. Using a genetic algorithm, we found sets of genes whose expression in the cell lines resembled the morphological distances obtained for each network. We then performed Gene Ontology analyses to assess the enriched cellular compartments in which the proteins encoded by these genes are present (Figure 4D). Interestingly, we observed that the genes whose expression was similar to the distances obtained by the 15px network are enriched mainly in the nucleus and also in the plasma membrane. The genes with similarity to the 30px and 45px network have more presence in cytosolic terms, while genes resembling the 60px network results are mainly enriched in plasma membrane and extracellular regions. This result suggests that morphological distances from different cellular compartments resemble molecular changes associated with these specific locations.

**Figure 4.**
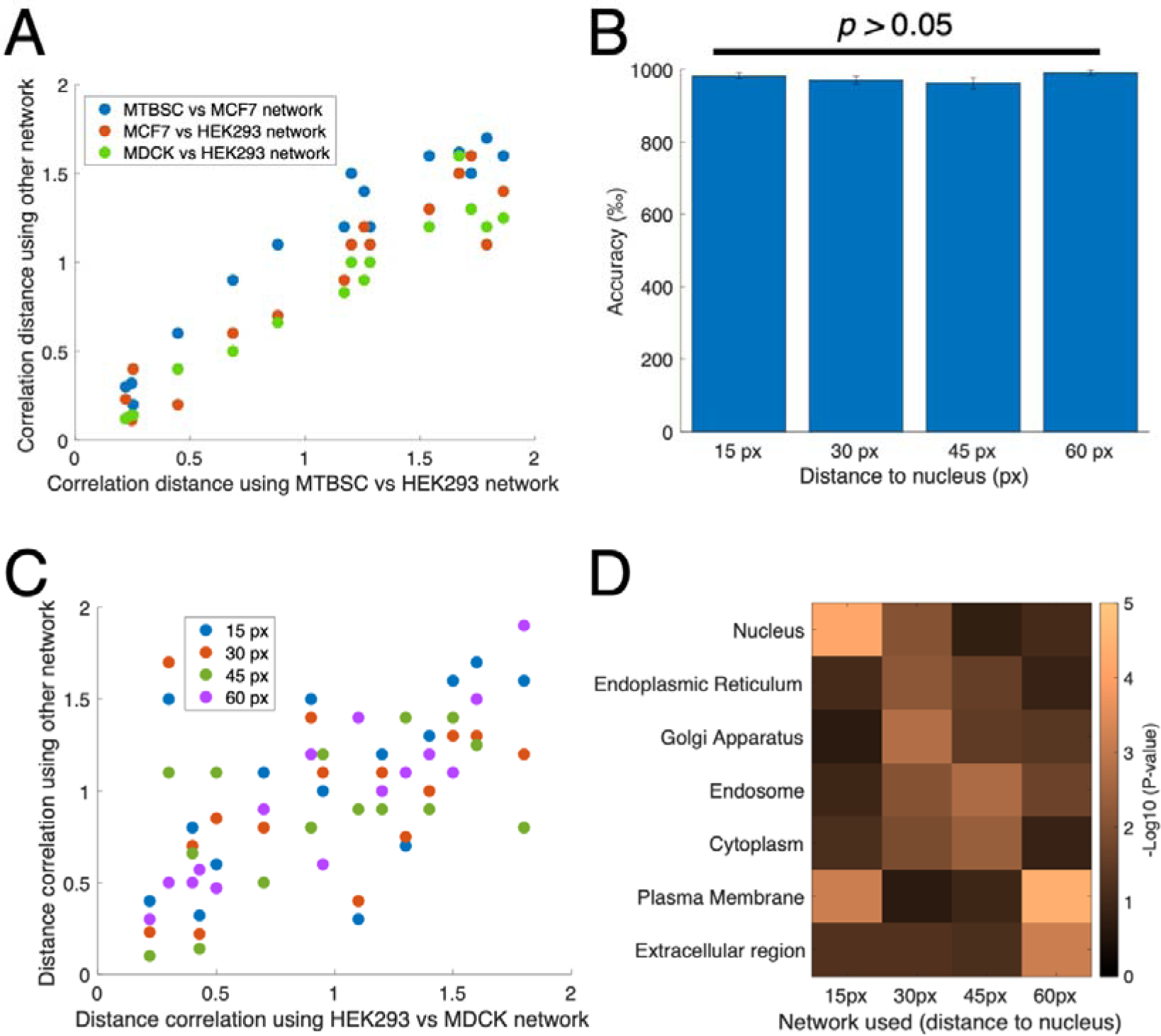
Morphological distances resemble molecular profiles. (A) Comparison between the morphological distances (using correlation distance) of cell types using the MTBSC vs HEK293 network (X-axis) and other networks specified in the legend (Y-axis). (B) Accuracy in the detection of cell lines using subcellular networks (15px, 30px, 45 px and 60 px). (C) Comparison between the morphological distances (using correlation distance) of cell types using the HEK293 vs MDCK network (X-axis) and other networks specified in the legend (Y-axis). (D) Heatmap representing the p-values (as -log10) of different Gene Ontology (Cellular Component) terms that are obtained from sets of genes identified to optimize the similarity between morphological distances (of subcellular networks) and transcriptional differences of six human cell lines (HEK293, MCF7, MDA-MB-231, HEPG2, HL-60, and SH5-SY5Y).

Our data powerfully demonstrated the ability of subcellular regions to determine cellular and molecular identity, but we wanted to support this idea with another test. We used short pharmacological treatments to change cellular morphology in a controlled manner. Specifically, we treated HEK293 cells for one hour with etoposide, an inhibitor of Topoisomerase II and producer of DNA damage (28), and nocodazole, a disruptor of the cytoskeleton (29). These cells were imaged, and we used the 15px, 30px, 45px, and 60px networks to observe the morphological changes produced by the drugs (Figure 5). With the 15px network, only the etoposide treatment produced a strong morphological change in HEK293 cells, consistent with the effect of this drug on the nucleus (Figure 5A). In the case of the 30px and 45px networks, the nocodazole treatment caused a major change in the cytosolic morphological features (Figure 5B and C). Finally, we did not observe significant morphological changes in the membrane features (60px network) of the HEK293 cells treated with either etoposide or nocodazole (Figure 5D). In conclusion, we validated the importance of morphological features for cellular identity.

**Figure 5.**
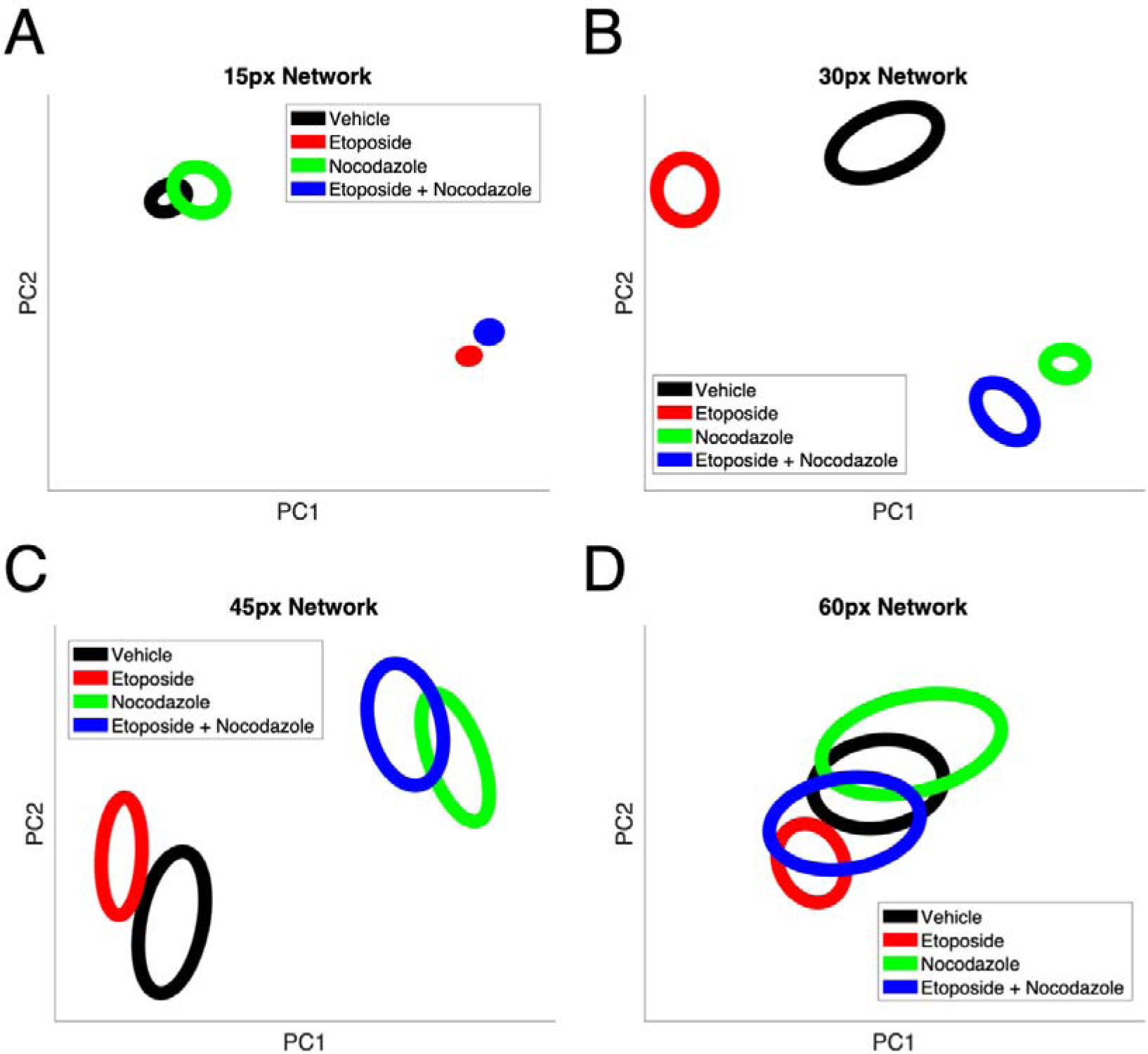
Morphological distances obtained after drug treatments. (A) Visualization of morphological distances (as in Figure 3D) using the 15 px network between HEK293 cells treated for 1 hour with vehicle, etoposide, nocodazole, and etoposide + nocodazole. (B) Same as A, but using 30 px network. (C) Same as A, but using 45 px network. (D) Same as A, but using 60 px network.

## Discussion

In this study, we demonstrate how subcellular regions contain sufficient information to distinguish and even quantify differences between specific cell types. We employed a straightforward CNN to model these features. It is highly likely that increasing the complexity of the network layers or changing its architecture would enhance detection accuracy and enable simultaneous classification of multiple cell types (30). Additionally, we utilized a simple method to extract a feature vector from the neural network. However, it’s worth noting that there exist specialized network architectures designed to optimize the use of feature vectors for estimating distances between categories (31). Nevertheless, our primary objective in this manuscript was to develop a broadly applicable method related to the information contained within subcellular regions, rather than fine-tuning conditions for each individual classification or measurement presented in the data.

An advantage of such a general approach lies in its adaptability. With simple modifications, we have successfully performed multiple tasks that could prove useful for scientists. For instance, estimating the number of distinct cell types within a co-culture holds interest across various areas of stem cell biology and cell differentiation processes (10, 32). It can also be applied in functional assays, such as wound healing (33). Another intriguing future feature would be real-time cell type detection directly under the microscope, leveraging marker-free subcellular morphology analysis. Adapting this methodology to microscopy software could significantly expand the toolkit available in cell biology (34). Furthermore, we anticipate that scientists will envision other utilities for subcellular regions within biomedical contexts. The potential utilization of networks that do not even include the original cell types targeted for quantification (as demonstrated in Figure 4A) further broadens the possibilities for studying complex or dynamic cellular samples.

Here we established a clear correlation between morphological distances and molecular distances. Specifically, transcriptomic and morphological profiles exhibit strong correlation (as depicted in Figure 4D), supporting the idea of cellular identity. Morphological features distinguishing between different nuclear, cytoplasmic or membrane regions directly indicate changes in the expression of nuclear, cytoplasmic or membrane genes, respectively. One remaining question is whether this approach can identify novel genes related to nuclear structure, for example. Future experiments could explore this idea by modulating the expression of potential target genes and observing resulting changes in the morphology of specific cellular compartments. Additionally, a network trained to identify malignant (tumoral) features within cells might help identify genes associated with malignancy, potentially leading to the discovery of novel targets in Oncology.

In a similar way, our latest results in the manuscript (Figure 5), support the correlation between cell morphology and molecular profile and also open the possibility of adapting this approach to drug screening and discovery methods (35). A library screening followed by cell imaging and analysis with the 15px, 30px, 45px, and 60px networks would identify drugs that specifically affect different cellular compartments. Similarly, the use of a network that quantifies malignant features would identify drugs with potential anti-tumoral effects. The only difference between the two analyses is the use of a different network, so the cell images obtained after a drug screening could be reused for different purposes. This is very important for perturbation screenings, which are a relevant pillar in current biomedical and pharmaceutical scientific fields.

## Materials and Methods

### Cell culture

All the cell lines were cultured at 37°C and 5% CO_2_, and media used followed the recommendations of ATCC. Cell lines were purchased by ATCC, except in the cases of normal human dermal fibroblasts from adults (NHDFs, Lonza), human papillary and reticular dermal fibroblasts (Tebu-Bio). Mouse tracheal basal stem cells (MTBSCs) were extracted and cultured as we previously published (26). Briefly, MTBSCs were isolated from wild-type C57BL/6J adult mice that were sacrificed and tracheas were dissected from the bronchial main to the larynx and collected in cold Ham-F12 containing penicillin and streptomycin (Gibco). After elimination of vascular, fatty tissues, and muscle, clean tracheas were excised longitudinally and then incubated in Ham’s F-12 pen-strep with 1.5 mg/mL of pronase (Roche Molecular Biochemicals) for 16 hours at 4°C. Processed tracheas in 10% fetal bovine serum were discarded, and the isolated cells were collected by centrifugation. After pancreatic DNAse I (Sigma) incubation, cells were collected again by centrifugation. Finally, MTBSCs were seeded within PneumaCult-Ex Plus complete medium (StemCell) in primary tissue culture plates (Corning) for 4 hours in 5% CO2 at 37°C to remove fibroblasts. Supernatant was collected, and MTBSCs were seeded in type I rat tail collagen (Gibco)-coated plates for routine culture and analysis after 8 and 12 passages. In the case of human fibroblast clones, we used a previously established protocol (36). Briefly, skin biopsies were cultured in gelatin (Sigma)-coated plates for around 2 weeks, when fibroblasts sprouted from their edges. Then, cells were split for four passages using trypsin (Gibco), and different clones were imaged and analyzed.

### Cell Imaging

For all the experiments presented in the manuscript, we used an EVOS FLoid microscope (ThermoFisher Scientific) with its 20x objective. Using a different microscope with the same objective produced similar results (data not shown). Raw TIFF grayscale images (20-40 for the training of a specific label in a network and 5-10 for testing) were sent to MATLAB to subsequent analyses.

### Pre-processing of images

We developed a MATLAB function that extracted ROIs of determined size (24×24 and 32×32 pixels) from the original images. This function also converted the grayscale image to black and white using Otsu’s method, and then quantified the percentage of area in the ROI that was above this threshold. In order to remove ROIs with low cell content, only the ROIs with 70% or above of their area considered signal were maintained in the following step.

### Design of a convolutional neural network (CNN) for cell classification

We used the processed ROIs (40,000 to 100,000, the same number for every cell type) as input of different CNNs following a standard scheme proposed by MATLAB. It consisted on twenty seven layers (the input layer, six blocks of 2D convolutional, batch normalization, ReLu and max-pooling layers and finally a fully connected layer followed by a softmax layer) that were trained at an initial learn rate of 0,00001 during 100 epochs in an NVIDIA GPU Titan X (kindly provided by NVIDIA).

### Quantification of accuracy in the detection of cell types

A testing set of ROIs were used for prediction using the trained CNN, and accuracy was directly obtained as permille. For *in situ* prediction, we developed a MATLAB function that used running ROIs in a single image for pre-processing and prediction of cell type. As the running ROIs were overlapped, we were able to determine the predictions at a single pixel resolution if a specific cell type was predicted to be in more than 50% of the times. The determination of accuracy in this case was done using human validation and comparison with a live cell marker (a specific cell type was pre-treated with a red fluorescent marker, Abcam). Specifically, unbiased humans determined the contours of a cell type within a white light image, and accuracy of the prediction was obtained by dividing the sum of true positives and true negatives by the total number of pixels. As comparison, the signal obtained in the red channel was converted into black and white using Otsu’s method and then accuracy was estimated as with the prediction.

### Estimation of cell number

First, we established co-cultures by plating specific numbers of different cell types measured by a cell counter (TC10, BioRad). After 3 hours, medium was changed and then imaged. For the quantification of the percentages of different cell types within a co-culture, we proceed as in the *in situ* prediction with minor modifications. Specifically, we manually estimated the pixel size of each cell type and then we normalized the pixel area of each cell type by its estimated cell size. Then, we fitted the obtained vs. expected percentages using a linear model. In the case of detachment, MDCK cells were imaged during trypsin treatment. We also generated a CNN using as input the initial (time = 0 seconds) cells and final (time = 420 seconds) cells.

### RNA extraction and gene expression

Total RNA from cells was obtained using Purelink kit (ThermoFisher Scientific); then 400 ng of RNA were reverse transcribed to cDNA (High Capacity cDNA Reverse Transcription, ThermoFisher Scientific). Quantification of expression was done by qPCR with specific oligos using GAPDH as normalization control.

### Morphological distance between cell types

We used the previously described CNN for estimation of morphological distances between cell types. Specifically, we used the last max pooling layer in order to have a feature vector (embedding) of 420 variables that can be used for distance estimation. Then, we are able to extract this feature vector from each ROI that is going to be analyzed. For visualization, we used PCA in the ROI population for reduction to only two dimensions and then we utilized the eigenvectors for estimation of an ellipse, as we did in previous works (37). For distance estimation between cell types, we used the median vector of each cell type and then we quantified the correlation distance (1 – correlation), obtaining a distance between 0 and 2 for every pairwise comparison between cell types.

### Correlation between transcriptional profiles and morphological profiles

We first established four different CNNs whose raw input images were the same but differed in the pre-processing. The pre-processing was done as previously defined (using a contrast detection threshold for defining ROIs) but with a modification. In this case, we used Hoechst 33342 (ThermoFisher Scientific) to label nuclei of living cells, and an *ad hoc* MATLAB function allowed us to estimate the relative distance between the preprocessed ROI and the centroid of the nearest nucleus. Thus, we could assign ROIs that were closest to the nucleus (less than 15 px) and more distant (from 15 px to 30 px; from 30 px to 45 px, and finally from 45 px to 60 px). Finally, we can use these different ROIs as input of the CNN that learns to distinguish different features that are near to the nucleus in HEK293 and MDCK, for example. For comparison with transcriptional profiles, we retrieved the transcriptome of HEK293, MCF7, MDA-MB-231, HEPG2, HL-60, and SH5-SY5Y cell lines from CCLE (27). We also quantified the morphological distance between these cell lines in a pairwise comparison as previously described using the four subcellular CNNs but using cosine distance. Then, we optimized the genes whose expression was explaining the morphological distances between cell lines in each network using a Genetic algorithm. Specifically, we randomly established a population of 10,000 individuals with 300 genes each from the transcriptome. We estimated the pairwise distance between cell types from each individual using cosine distance of their gene expression. Then, we selected the top 50 individuals (with minimal difference to morphological distance) as parentals for the next generation (10,000 individuals), and simulated a cross between two parentals with a random mutation rate of 0 to 120 genes. After 30 generations, we selected the top individual that minimized the difference between transcriptional profiles and morphological distances. Then, the set of 300 genes was subjected to Gene Ontology, and the values for cellular compartment terms were analyzed in the different subcellular networks.

### Etoposide and Nocodazole treatments

We treated HEK293 cells with dimethylsulfoxide (DMSO, vehicle), 20 uM etoposide, 50 nM nocodazole or the combined etoposide plus nocodazole treatment for 1 hour. After imaging the cells, we performed distance estimation of their ROIs using the 15 px, 30 px, 45 px and 60 px networks, and ellipse visualization as described before.

### Statistics, data and code availability

We performed three replicates of each test presented in the manuscript, and in some cases we present a single experiment for visualization purposes. If not specified, paired T-tests are used for statistical comparison. In the case of coefficients of correlation, p-values are computed by transforming the correlation to create a t statistic having n-2 degrees of freedom, where n is the number of individuals. Raw data as well as MATLAB code is available in Figshare.

## Acknowledgements

This paper was supported by the Spanish Ministry of Science through the projects TED2021-130036B-I00 and PID2020-117467RB-I00 (to SMN and ACR), and PID2021-126905NB-I00 and TED2021-130560B-I00 (to JMCG).

